# Qualitative assessment of cryo-EM map reconstruction

**DOI:** 10.1101/2022.12.31.521834

**Authors:** Nidamarthi. H. V. Kutumbarao

## Abstract

With the rapid advancement of Cryo-EM technique in deciphering structural features of biological molecules, it is a necessity to develop and explore different parameters which help us in critical evaluation of different aspects of reconstruction. In this age of ever-increasing automation and number of particles used in reconstruction to achieve high resolution maps, assessment parameters help in validation of reconstruction process. Here, we have employed one such quality parameter which quantitatively assess the presence of non-particles in the reconstructed map.

## Introduction

Cryo-EM is one of the most popular techniques to study biomacromolecules, especially those with higher molecular weight, and complexes. With the rapid development in detector’s technology and computational resources, single particle analysis, along with crystallography, has become an integral tool in deciphering the structure-function relationship of macromolecules ^1,2^.

The output of a cryo-EM experiment is an electric potential map. This 3D density object is reconstructed from 2D projections of a protein^3,4^. The basic approach used in 3D reconstruction is an extension of 2D reconstruction from 1D projections. The reconstruction of map from a set of projections has been implemented for a long time^5^. In this process many projections are obtained as single images and later combined after derivation of orientation parameters, where each projection is a footprint of many orientations of the macromolecule under the electron beam.

Thus, single particle Cryo-EM is an averaging process over a large set of 2D images, to obtain a 3D map via computational techniques. The resolution of the resulting highly averaged map is not uniform over all regions. As a matter of fact, the concept of resolution in cryo-EM maps is not so much to do with the resolving power of the microscope or the distinguishability between intensity maximas, rather, resolution is portrayed as the maximum spatial frequency at which the information content can be considered reliable. To represent the concept objectively, the resolution is generally assessed using a correlation called Fourier shell correlation (FSC), where the information agreement at different Fourier shells between two halves of reconstructed map is calculated^6,7^. The said agreement can also be interpreted as a self-consistency test of the resultant maps^8^. In exploring the correlation, it has become general practice to interpret the curves at a certain threshold^9,10^. The magnitude of the threshold value employed is a controversial and a frequent topic for discussion^11,12^ and interpretation^13^.

In retrospect, FSC is one of the main and frequently used assessment methods to explain the quality of reconstruction, and also a namesake for resolution. FSC is a correlation between two halves of the reconstructed map, and hence, might not, in fact represent the validity and valuation of the reconstruction procedure^14^. There are few pressing concerns related to the concept of reconstruction, like the use of projection data with non-uniform SSNR and non-uniform or incomplete set of projections^15^. Furthermore, there are other significant points to be addressed, such as, the proportion of meaningful projections (particles) used for reconstruction, the particles chosen for reconstruction cover the complete distributions of projections possible and finally, how can the process of appropriate alignment of particles be quantified and the discrepancies in the final map be evaluated^16^.

In crystallography, the diffraction experiment can itself provide relationships between different two-dimensional gratings so that orientations are a *prior* experimental information and the intensities can be judged for symmetry relation among different measures. There are many residual factors and parameters in crystallography which are employed to measure the quality of a dataset, and up until recently the exploration for such measures was a thriving research field^17^. Quite differently, cryo-EM maps and their qualitative interpretation are only partially recovered from structural features (real space domain), and yet the whole idea and backbone of reconstruction methods is rooted in Fourier space. Thus, the Fourier space is an appropriate tool for a proper qualitative assessment of cryo-EM reconstructions. In the Fourier domain, phases are considered to impart more structural information than magnitudes ^18^.

A frame work has been employed in this study to understand the information flow with respect to phase values in cryo-EM maps. Such a validation of reconstructions is an indication that the electron scattering information obtained as 2D images has been positively transformed into the appropriate 3D volume. Thus, the structure and its functional implications can be interpreted properly. If such validation is not reached, maps tend only to explain high resolution surface information, thus contradicting the concept of structure analysis. This paper introduces and explains a new procedure to validate the reconstruction of maps from cryo-EM experiments.

## Method

Present analysis rests on the assumption that, in Fourier domain of a density map, phase values tend to have a relation with its adjacent values. Such a relation might manifest into statistical features, especially when the density values are from structural data. Such a statistical measure can be used to distinguish maps which are generated from non-particles or noise images from that of structural features.

Maps are first preprocessed to create a mask outside the region of interest. Tight masking is performed using the skimage’s morphology module and further the noise dust present surrounding the region of interest is removed^19^. The author provided contour value is used to calculate the threshold values for creating a mask. The analysis is carried out in two steps, firstly, Fourier transform is performed on density values from which phase values are extracted upon which Phase coherence (PC) are calculated. Both these steps are performed along diagonal axis (depth) of 3d array, the calculation can be performed along any of 3d array. (Figure 1). This step can be perceived as a dimensionality reduction method, where we obtain a 2-dimensional array of PC values which represents a 3D map.

**Figure 1:**
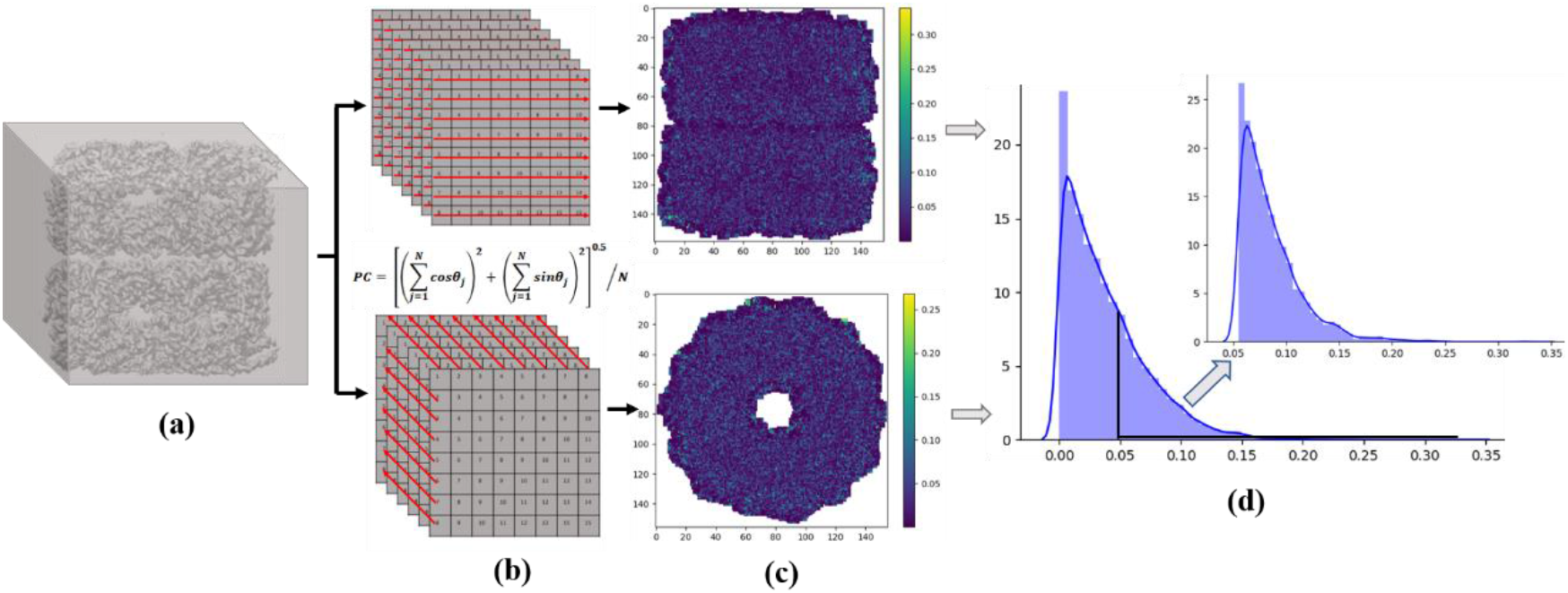
Overview of the method (a) 3D image of a cryo-EM map (b) calculating PC values along each of the array (along the diagonal or horizontal axis) (c) pictorial representation of PC values obtained from respective axis (d) the distribution of PC values calculated from any of the axis, the inset plot shows distribution of 75^th^ percentile PC values.

The Phase coherence is a way to evaluate variation of phase values and its mathematical form is given below,

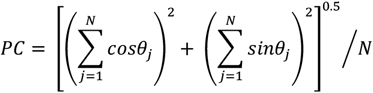

where *θ*_*j*_ is phase value obtained from each density point
The importance of PC values comes from the drawback pertaining to traditional variance and standard deviations, which are not appropriate to understand the variability of angular data (phase values) because of the non-compatibility with a break at 360º^20–22^.

PC is inversely proportional to phase variability and takes values between 0 and 1. When PC is minimum, the variability of phases is maximum and vice versa. This implies, the phase variability of structured data will be higher when compared to maps generated from noise (where phase variability is less). A characteristic distribution of PC values (which are high in magnitude) can be observed when calculated from various simulated maps. Further, the data from 75^th^ percentile are used for the analysis, where, the shape of the histogram pertaining to data between the 75^th^ and 100^th^ percentile (inset graph in fig 1) is quantified using the skewness and kurtosis test values. These test values indicate the presence of non-particles present in the dataset considered for reconstruction, and thus can help us in qualitative assessment of the reconstruction process.

## Results and Discussion

Phase analysis is carried out along diagonal axis of density points in a map resulting in the reduction of 3-dimension map to 2-dimensional arrays of PC values. Each value from this obtained 2D array indicate phase variability along an array of density points in a map. The PC values are used for statistically descriptive judgment of the reconstructed maps from projections. In an ideal situation there should be gradual fall (Fig 2) in the high PC values (75^th^ percentile data) indicating the reduction of less phase variability regions which signifies lower representation of non-structured particles (projections) considered for reconstruction. Such a behavior (of PC values) can be observed in simulated maps (Fig 2), crystallographic map and in good reconstructed cryo-EM maps (Fig 3).

**Figure 2:**
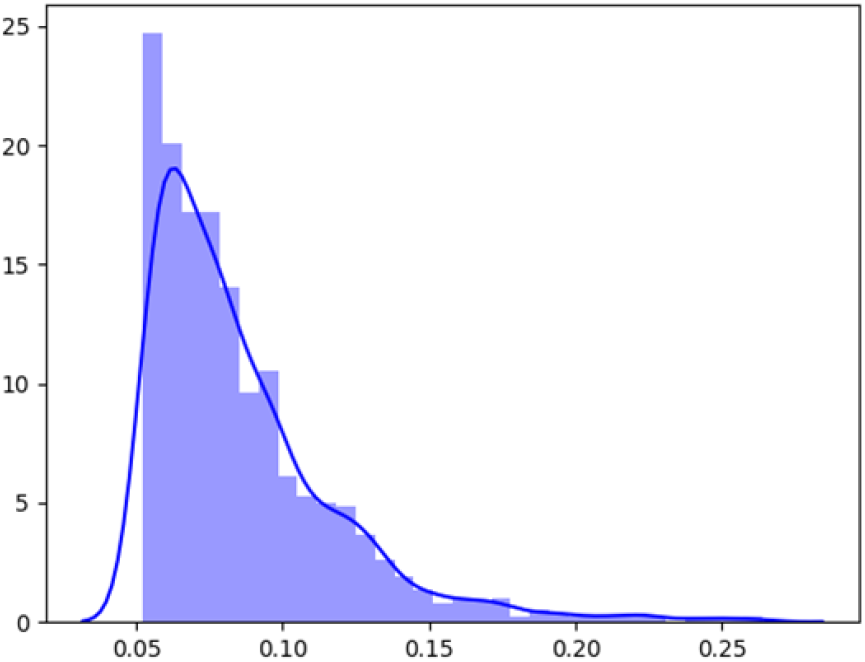
The distribution of high PC values from a simulated map

**Figure 3:**
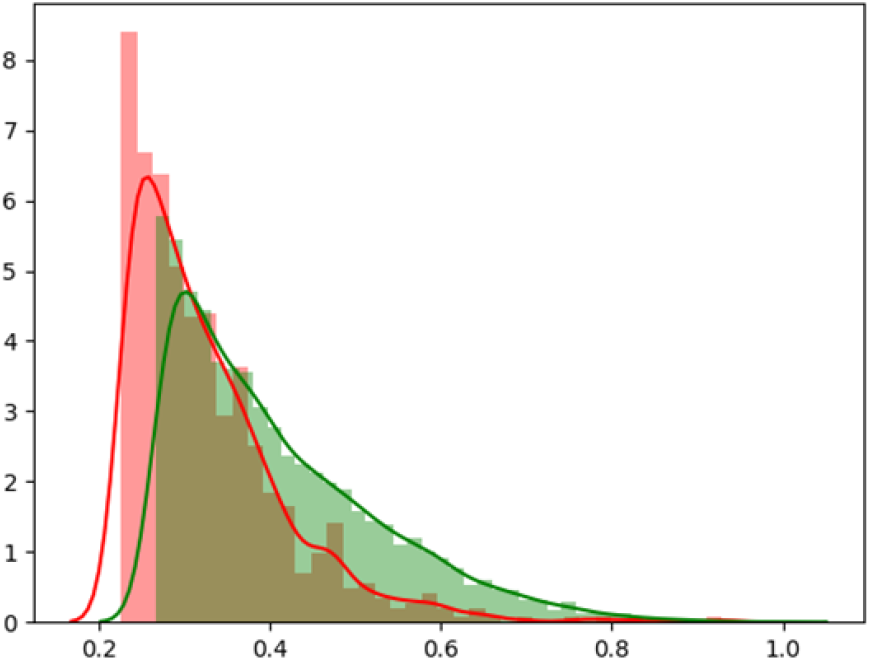
The distribution of high PC values from a crystallographic map (green line) and cryo-EM map (red line)

The Z-score values of skewness and kurtosis tests are used to represent the characteristic distribution of PC values. The scatter plot of Z-scores (kurtosis tests vs Z-scores of skewness test) should bring out a characteristic slope in the first quadrant of a graph as observed when plot was derived from PC values calculated from ∼1000 simulated maps (wide spectrum of protein structures was considered) (Fig 4). On the other hand, for certain maps which were obtained from cryo-EM reconstructions (experimental) we can observe a marked difference in the Z-score values. To further understand the implications of the difference in the shape of distribution of high PC values, we have carried out the analysis on maps from GroEL derived from simulations, crystallography, and cryo-EM processes. The simulated maps (with different resolutions) were generated using 1sx3. In figure 5, blue dots indicate simulated data, green dots represent maps from crystallography experiment and red dots from cryo-EM experiment. We can observe points spread into the third and fourth quadrant of the graph, such values indicate discrepancies in the maps generated. These values signify a marked change in the distribution of high PC values (less phase variability). Such a contrasting distribution, points out the disparity in the reconstruction process. The disparity might arise either from inclusion of non-particle images or particles which have high noise content. For convenience, we can label the lower two quadrants of scatter graph (third and fourth) as disallowed regions with respect to the reconstruction process. Additionally, we can also use the 75^th^ percentile value as a magnitude cut-off to represent a good reconstruction, where, for simulated maps and for experimental maps (where the fall off of PC values are gradual and reconstruction is without disparity), the 75^th^ percentile value is lower when compared to the maps which have noisy particles crept into the reconstruction process.

**Figure 4:**
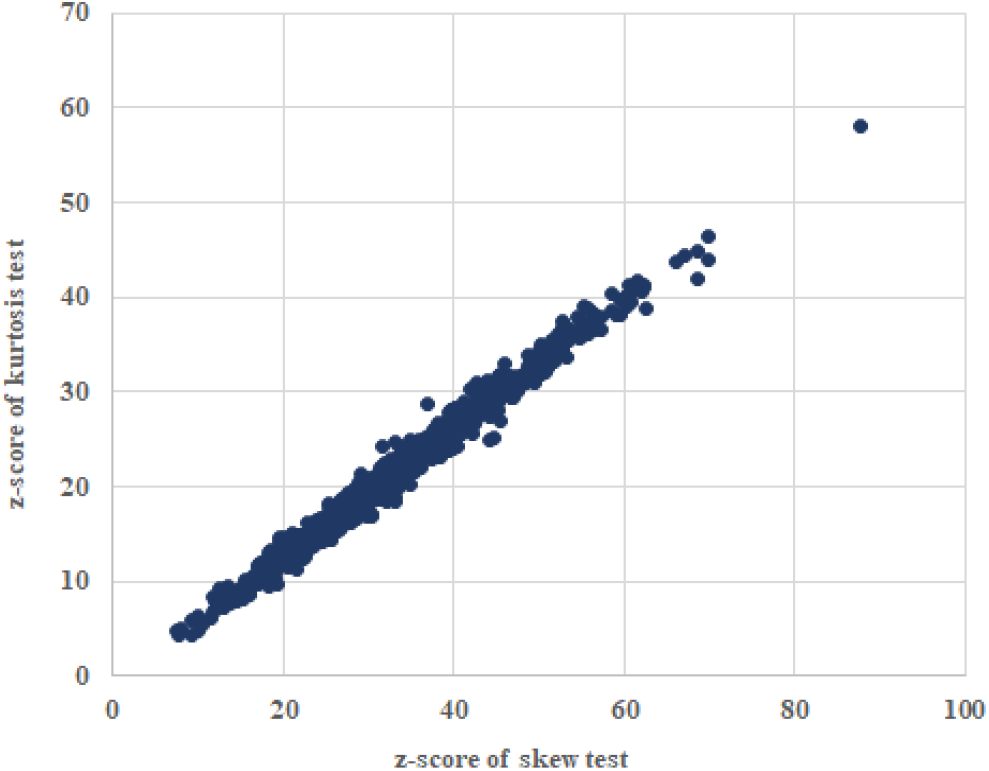
Scatter plot of Z-score of skew test and kurtosis test from simulated maps

**Figure 5:**
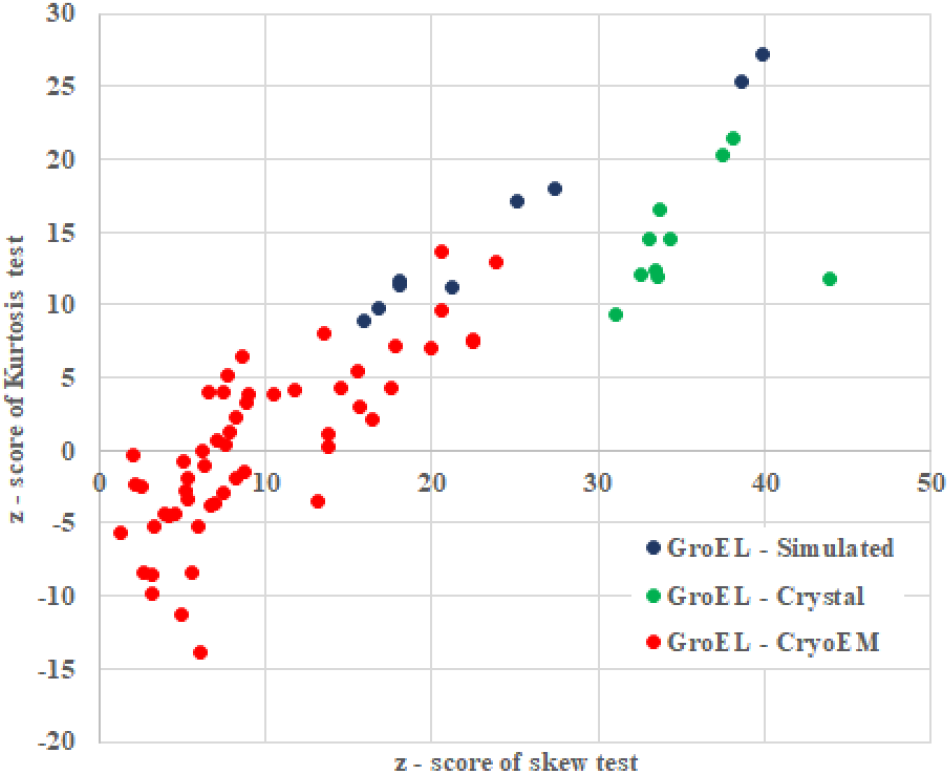
Scatter plot of Z-score of skew test and kurtosis test, GroEL-simulated maps (blue), crystal maps (green) and cryo-EM maps (red)

When the analysis was extended to various other cryo-EM maps (84 maps), values are spread into the fourth quadrant (red) can be observed (Fig 6). however, certain high-resolution maps (green) chosen for the study, show values in the first quadrant like those observed in maps from simulated and crystallographic studies (Fig 6). There has been a surge of structural studies from both crystallographic and single particle analysis of proteins from the COVID-19 virus. Around 350 maps corresponding to COVID-19, deposited in EMDB, were analyzed for discrepancies. The graph shows data points corresponding to maps which fall in the allowed first quadrant region, but there are quite significant number of maps which are observed in the disallowed region (Fig 7). Such maps indicate the presence of noisy non-particles in the final reconstructed map. It is crucial to evaluate the projections during the reconstruction process for their presence will affect the quality of reconstruction and the information which the protein under study has to offer.

**Figure 6:**
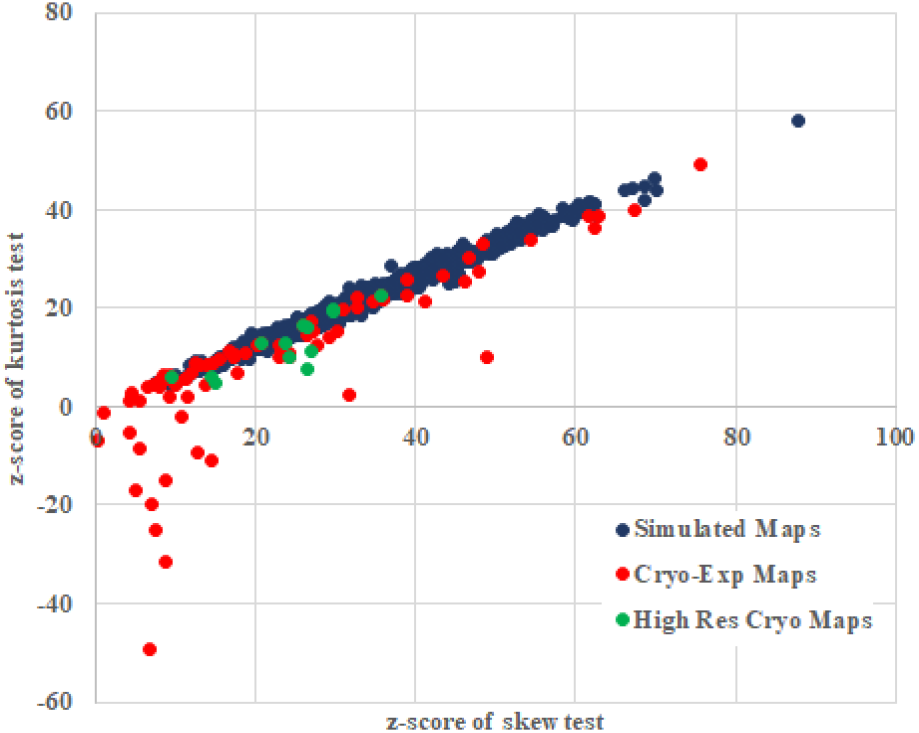
Scatter plot of Z-score of skew test and kurtosis test from simulated maps (blue), cryo-EM maps (red), High resolution cryo-EM maps (green)

**Figure 7:**
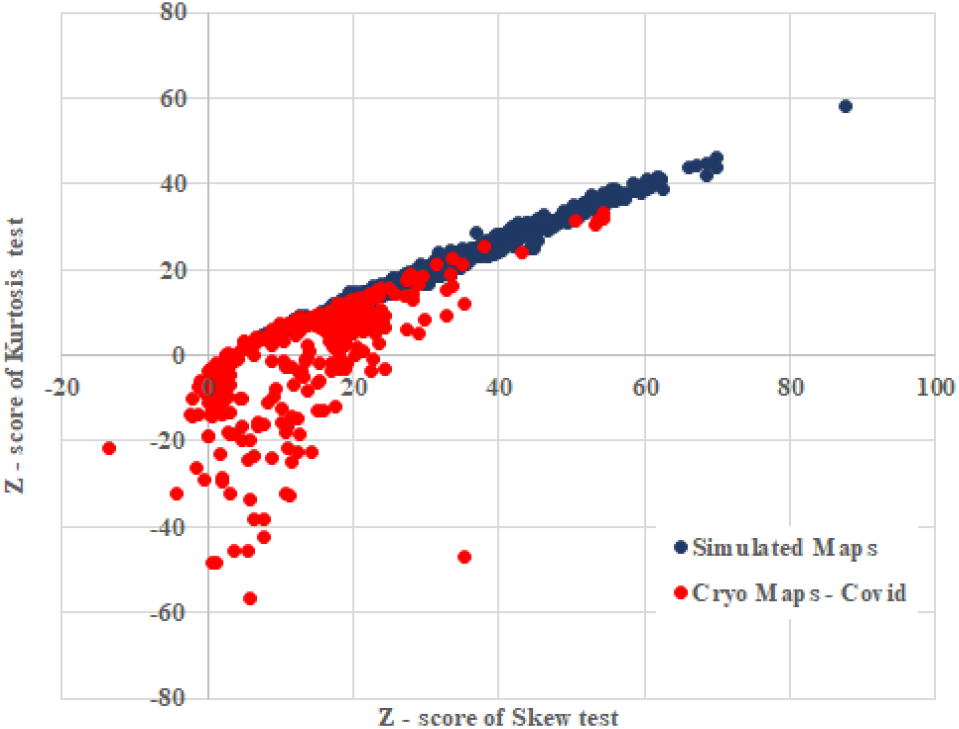
Scatter plot of Z-score of skew test and kurtosis test simulated maps (blue), maps belonging to COVID-19 virus (red)

This promising result led us to examine maps which were subjected to scrutiny in the past for their discrepancies^23,24^. In those studies, a plethora of non-particles were suspected to be considered during the reconstruction process^25–27^. It can be observed from Figure 8 that the Z-scores of these two maps suggest the presence of discrepancy in line with the previous inferences. The score values can be attributed to marked differences in the distribution of higher PC values compared to that of simulated and crystal maps (Fig 9). Additionally, the percentile cutoff values for both the maps are higher (∼0.4) when compared to that of simulated and other cryo-EM map with proper reconstruction. Such deviation in the distribution of higher PC values indicate presence of increased lower phase variability regions in a cryo-EM map.

**Figure 8:**
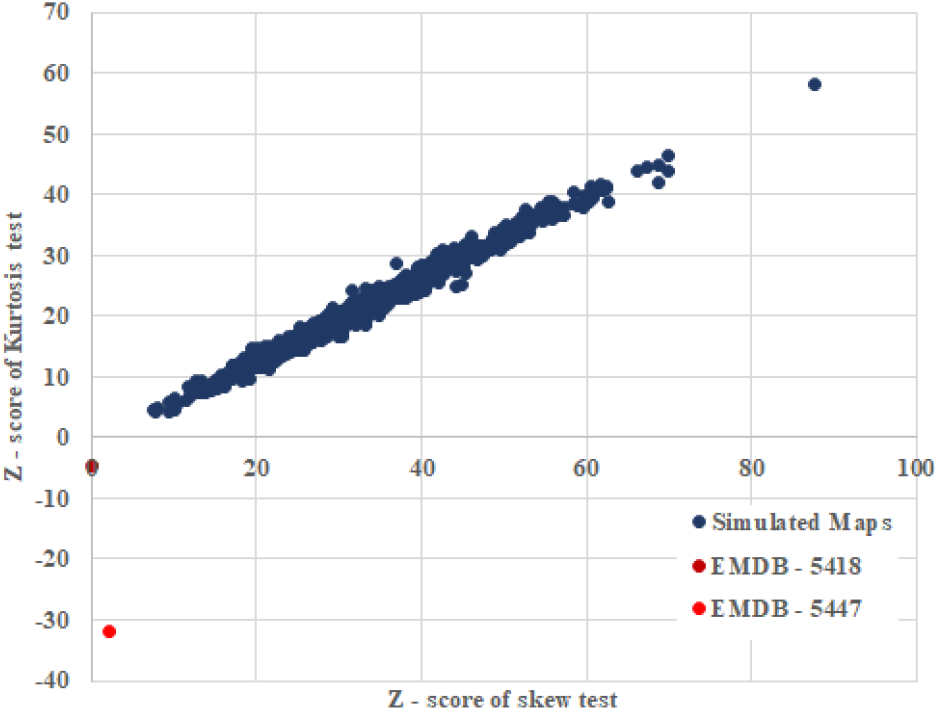
Scatter plot of Z-score of skew test and kurtosis test from simulated maps (blue), EMDB-5418 (maroon), EMDB-5447 (red)

**Figure 9:**
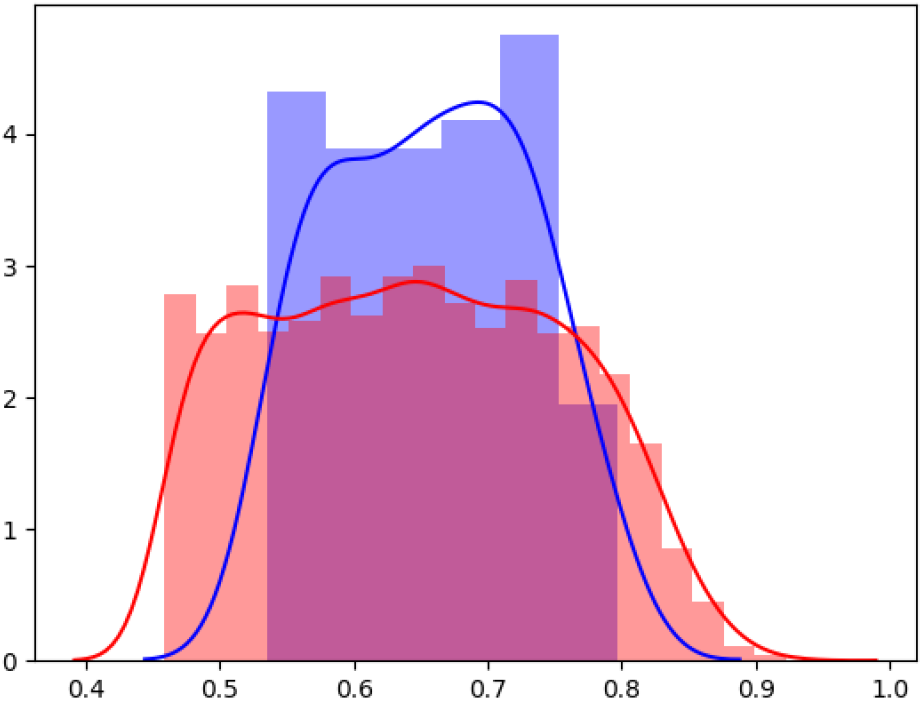
The distribution of high PC values of EMDB-5418 map (blue) and EMDB-5447 map (red)

To understand weather the Z-scores are influenced by map resolution, a scatter plot of the scores along with the resolution was produced. It is observed that in the disallowed region, there are more high-resolution maps with discrepancies than low-resolution maps, indicating that the Z-scores of maps falling into the third and fourth quadrant (disallowed regions) are not dependent on resolution (Fig 10).

**Figure 10:**
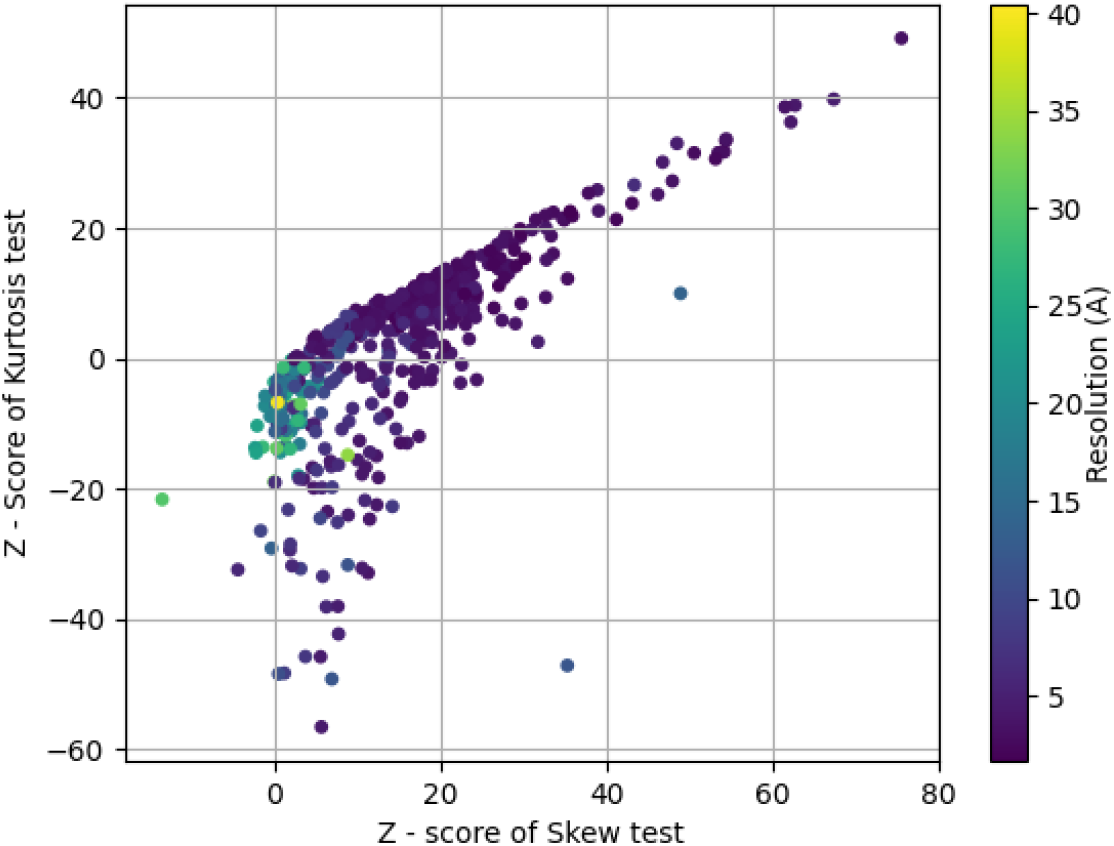
Scatter plot of Z-score of skew test and kurtosis test from cryo-EM maps with resolution.

## Conclusion

There are many studies elaborating the importance of meaningful particle picking, judging particles and assessment of particles by visual inspection, but the automation procedures and assumptions from an experiment might lead the reconstruction process to tolerate non-particles. Furthermore, the reconstruction method and performance of many algorithms have grown larger and larger so to facilitate convincing reconstructions from partially correct particles. The sound example of Einstein from noise^28^ has been demonstrated and also used as an argument to the assessment of the final reconstruction. Despite the continued development in algorithms and machine capabilities, qualitative measures to evaluate 3D reconstructions has been restricted to FSC and tilt pair parameter. To this end, a preliminary descriptive analysis such as the one explained above and more robust analysis on phase variability can help us in the assessment of the reconstruction process and guide us in deciphering more meaningful information from the maps.

